# Cellular senescence dysregulates antiviral interferon responses in idiopathic pulmonary fibrosis

**DOI:** 10.64898/2026.04.20.719739

**Authors:** Jun-Wei B. Hughes, Yasmina Reisser, Franziska Hornung, Tyler A. U. Hilsabeck, Fiona Senchyna, Ana L. Coelho, Tsung-Che Ho, Kevin Schneider, David Furman, Cory M. Hogaboam, Claude Jourdan Le Saux, Pierre-Yves Desprez, Stefanie Deinhardt-Emmer

## Abstract

Patients with idiopathic pulmonary fibrosis (IPF) are highly vulnerable to respiratory virus infections, but the cellular mechanisms linking fibrotic remodeling to impaired local antiviral defense remain unclear. Here, we investigated how cellular senescence shapes the response of patient-derived healthy and IPF primary lung fibroblasts to influenza A virus (IAV) infection. Transcriptomic profiling identified infection as the driver of gene expression in both DNA damage-induced senescent healthy and IPF fibroblasts and revealed induction of canonical antiviral pathways in both cell states. However, senescent IPF fibroblasts adopted a distinct antiviral response state characterized by a broader set of uniquely induced genes and differential coordination of antiviral transcriptional networks. Functionally, senescence increased viral titers in healthy and IPF fibroblasts, while senescent IPF fibroblasts displayed an altered inflammatory response. Network analysis linked viral response- and cell cycle-associated modules specifically to the senescent healthy infected state, whereas these programs were weaker in senescent IPF fibroblasts. Transcription factor inference identified *IRF3* and *STAT1* as candidate regulators of this altered antiviral state in both senescent healthy and IPF fibroblasts. Consistent with the network and transcription factor analyses, siRNA-mediated depletion of *IRF3* or *STAT1* significantly reduced IFN-β secretion in senescent healthy fibroblasts, whereas IPF fibroblasts showed only milder effects, indicating a disease-specific dependence on these pathways for antiviral control. Together, these findings show that the combination of cellular senescence and fibrotic fibroblast identity creates a dysfunctional antiviral state that may help explain the high susceptibility of IPF patients to virus-associated acute exacerbations and disease worsening.

## 1 INTRODUCTION

Idiopathic pulmonary fibrosis (IPF) is a progressive, irreversible interstitial lung disease increasingly recognized as an age-associated disorder of the lung (King et al. 2011; Martinez et al. 2017; Maher et al. 2021). In current models of IPF pathogenesis, aging is no longer regarded merely as a demographic backdrop, but as a mechanistic driver of disease. Among the hallmarks of aging, cellular senescence has emerged as a key process contributing directly to aberrant tissue remodeling (Sucre & Plosa 2019; Álvarez et al. 2017; Schafer et al. 2017; Han et al. 2023). Experimental evidence further supports a causal role of senescent cells in fibrotic lung disease, as their accumulation is detectable in fibrotic tissue and their clearance can attenuate pulmonary fibrosis *in vivo* (Sucre & Plosa 2019; Álvarez et al. 2017; Schafer et al. 2017; Han et al. 2023).

Respiratory infections are clinically important in IPF, where they have been implicated not only as cofactors of disease progression but also as potential triggers of acute exacerbations, the most devastating complication of the disease (Molyneaux & Maher 2013; Mostafaei et al. 2021). Acute exacerbations of IPF are associated with very poor short-term outcomes, with reported mortality often exceeding 50% despite therapeutic intervention (Ryerson et al. 2015; Chen et al. 2024). This clinical vulnerability suggests that impaired local host defense is a relevant component of fibrotic lung disease.

Among respiratory pathogens, influenza A virus (IAV) represents a major cause of viral lower respiratory tract infection worldwide and thus provides a clinically and biologically relevant model with which to interrogate antiviral defense in the fibrotic lung. Respiratory viral infections, including influenza, have been implicated as potential triggers of acute exacerbations in IPF, further supporting the use of IAV as a model of infection-associated disease worsening (Sese et al. 2021; Kolb & Richeldi 2011).

Although epithelial injury remains central to prevailing concepts of fibrosis, fibroblasts are increasingly appreciated as immunologically competent structural cells that can sense danger signals, respond to inflammatory cues, and actively shape local innate defense programs (Cavagnero & Gallo 2022; Ghonim et al. 2023). Among the stromal compartments implicated in IPF, fibroblasts occupy a central position because they are both major effector cells of extracellular matrix deposition and active regulators of tissue remodeling (Álvarez et al. 2017; Schafer et al. 2017; Han et al. 2023). Various subtypes of fibroblasts isolated from IPF lungs display senescence-associated phenotypes characterized by altered survival programs, resistance to apoptosis, mitochondrial dysfunction, telomere abnormalities, and acquisition of a senescence-associated secretory phenotype (SASP), all of which are thought to reinforce profibrotic signaling within the diseased lung (Han et al. 2023; Schafer et al. 2017; Álvarez et al. 2017). However, despite substantial progress in defining fibroblast senescence as a driver of matrix remodeling, far less is known about how senescence shapes the capacity of lung fibroblasts to mount intrinsic responses to viral infection.

This question is particularly relevant in the context of IAV infection. Cell-intrinsic defense against IAV depends on rapid sensing of viral RNA and activation of interferon-centered transcriptional programs (Iwasaki & Pillai 2014; Opitz et al. 2007; Tolomeo et al. 2022; Talon et al. 2000). In lung cells, RNA Sensor RIG-I (*RIG-I*)-dependent signaling and Interferon Regulatory Factor 3 (*IRF3*) activation are critical for induction of type I interferons, whereas downstream transcription factors such as Signal Transducer and Activator of Transcription 1 (*STAT1*) orchestrate the antiviral interferon-stimulated gene response (Iwasaki & Pillai 2014; Opitz et al. 2007; Tolomeo et al. 2022; Talon et al. 2000). Aging, however, has been linked to impaired or dysregulated antiviral interferon responses, including defects in *RIG-I*-associated signaling, raising the possibility that age-related cellular states may uncouple viral sensing from effective effector function (Tolomeo et al. 2022; Molony et al. 2017). In the setting of influenza, fibroblasts are not passive bystanders but can adopt antiviral and pro-inflammatory states that influence tissue pathology and repair (Ghonim et al. 2023). Whether such rewiring also occurs in senescent IPF fibroblasts remains unresolved.

Addressing this question may provide mechanistic insight into why the aging fibrotic lung is particularly susceptible to infection-associated deterioration and acute worsening. We therefore investigated whether senescence in fibrotic fibroblasts amplifies antiviral signaling or instead drives a distinct host-response state in which viral sensing, interferon output, and functional control of infection become uncoupled.

## 2 RESULTS

### 2.1 Senescent healthy and IPF lung fibroblasts exposed to infection activate canonical viral response pathways with heterogeneity at the individual gene level

To understand functional differences between senescent healthy and IPF lung fibroblasts, we exposed healthy (3 donors) and IPF (4 donors) lung fibroblasts to 15 Grays γ-irradiation (IR), a well-established method that triggers senescence primarily through DNA double-strand breaks (DSBs) (Figure 1A) (Coppé et al. 2008; Schafer et al. 2017; Miller et al. 2025; Maus et al. 2023; Hernandez-Gonzalez et al. 2023). 10 days after IR, we exposed senescent healthy and IPF lung fibroblasts to influenza A virus (IAV) infection for 24 hours (h), a common infection globally serving as a model for the increased prevalence of viral infections in patients with IPF (Figure 1A). Principal component analysis revealed infection largely driving gene expression changes (Figure 1B). Differential expression analysis uncovered a larger proportion of upregulated than downregulated genes in both senescent healthy and IPF lung fibroblasts after infection (Figure 1C). Senescent IPF lung fibroblasts showed more upregulated genes than senescent healthy IPF lung fibroblasts with 160 unique, significantly upregulated genes in response to infection (Figure 1D). Shared, significant upregulated DEGs included viral response in both senescent healthy and IPF infected conditions, such as *RIGI* upregulation in both conditions (Figure 1E). Many upregulated genes in both senescent healthy and IPF lung fibroblasts after infection were related to viral response, as shown by Gene Ontology (GO) enrichment; however, there were some notable genes such as 2′-5′-oligoadenylate synthetase 1 and 2 (*OAS1* and *OAS2*) being significantly upregulated only in the senescent IPF infected condition (Figure 1F-G). Overall, senescent healthy and IPF lung fibroblasts exhibit similar responses to influenza infection at the global level, with notable differences at the level of individual genes.

**Figure 1.**
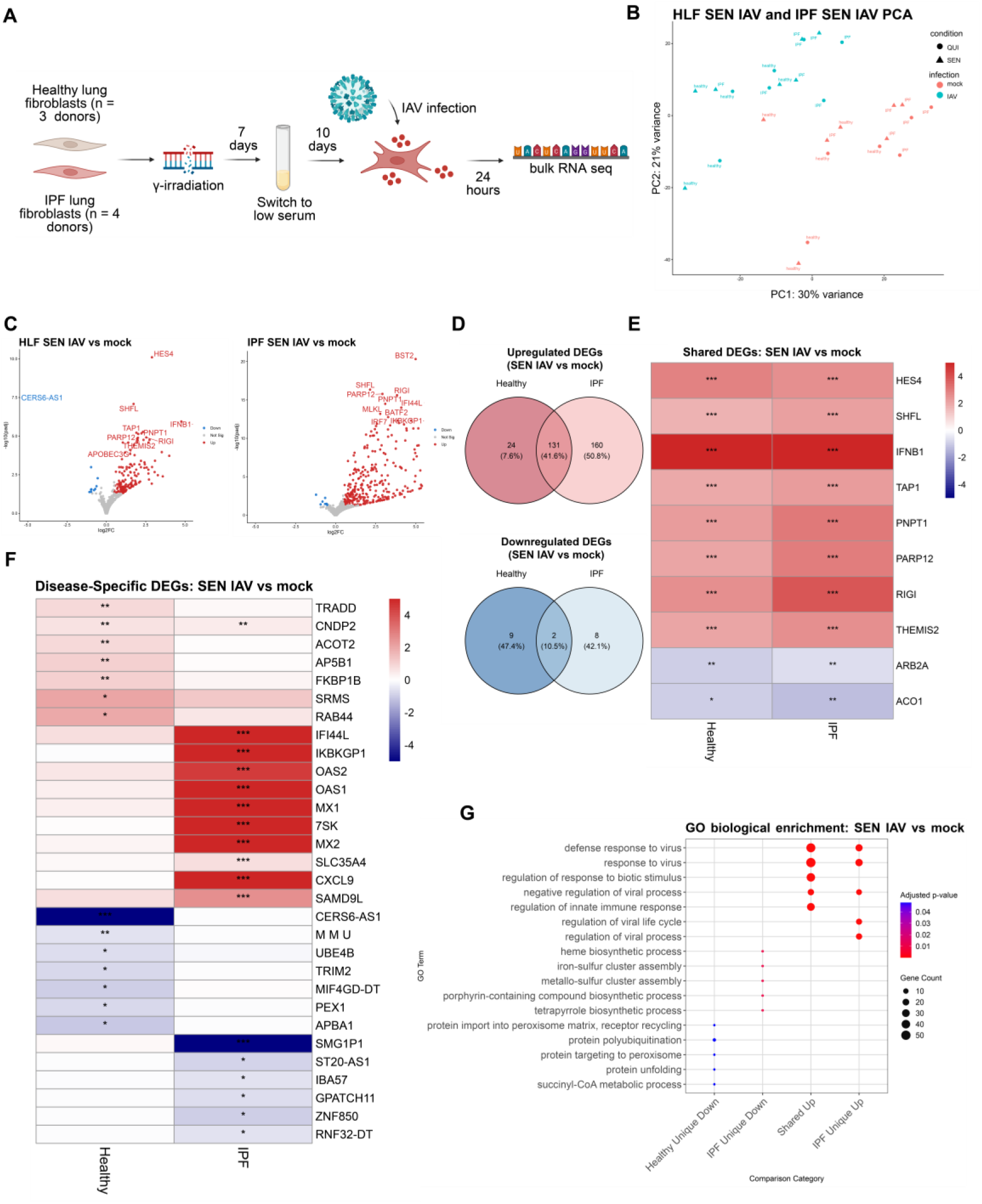
Senescent healthy and IPF lung fibroblasts exposed to infection activate canonical viral response pathways with heterogeneity at the individual gene level. (A) Schematic of 3 healthy (HLF) and 4 idiopathic pulmonary fibrosis (IPF) primary, human, lung fibroblasts, between ages 37-61, induced to senesce by γ-irradiation (IR; SEN). Quiescent (QUI) lung fibroblasts were changed to low serum (LS) 3 days prior to infection. Influenza A virus (IAV) infection was performed at 10-days after IR for 24 hours and RNA was collected for bulk RNA sequencing (RNA-seq) after the 24 hours infection period. (B) Principal component analysis (PCA) of QUI and SEN HLF and IPF lung fibroblasts with mock or IAV treatment. (C) Volcano plot of differentially expressed genes (DEGs) in HLF SEN IAV (left) vs IPF SEN IAV (right) lung fibroblasts. DEG cutoffs were defined by log2FC < -0.5 or > 0.5 and p-adjusted value < 0.05. Top 20 DEGs labeled and the volcano plot was capped at -5 to 5 log2FC. (D) Venn diagram of shared and unique upregulated (top) and downregulated (bottom) DEGs between HLF SEN IAV vs IPF SEN IAV lung fibroblasts. (E) Heatmap of shared top 10 upregulated and downregulated DEGs between HLF SEN IAV vs IPF SEN IAV lung fibroblasts. (F) Heatmap of unique top 10 upregulated and downregulated DEGs between HLF SEN IAV vs IPF SEN IAV lung fibroblasts. (G) Top 5 GO enrichment terms from HLF and IPF SEN IAV uniquely upregulated and downregulated genes and HLF and IPF SEN IAV shared upregulated and downregulated genes. Only 4 categories of GO enrichment shown as HLF and IPF SEN IAV shared downregulated genes and HLF SEN IAV uniquely upregulated genes did not enrich for GO terms.

### 2.2 Senescence in IPF fibroblasts results in decreased viral titers and altered cytokine production

To further investigate the impact of senescence on IPF fibroblasts in the context of viral infections, quiescent and senescent healthy and IPF fibroblasts were infected with IAV at a multiplicity of infection (MOI) of 1 for 24 h (Figure 2A). Specifically, changes in viral titers and cytokine release were analyzed. In both healthy and IPF conditions, senescent fibroblasts displayed an increased virus titer compared to the respective quiescent control fibroblasts (Figure 2B). However, the virus titer in senescent IPF fibroblasts was significantly lower than the virus titer in senescent healthy fibroblasts 24 h post infection (hpi). Interestingly, quiescent IPF fibroblasts showed a modest, non-significant decrease in viral titer relative to quiescent healthy fibroblasts. These findings imply a potential association between diminished viral replication and senescence in the IPF fibroblasts. A multiplex cytokine assay of the supernatant of mock and infected fibroblasts revealed a general trend to reduced pro-inflammatory cytokine production in infected quiescent and senescent IPF fibroblasts (Figure 2C). Specifically, IP-10 (*CXCL10*), critical in the inflammatory response to IAV, was significantly lower in both quiescent and senescent IPF fibroblasts, indicating a modified response of these cells to IAV infection.

**Figure 2:**
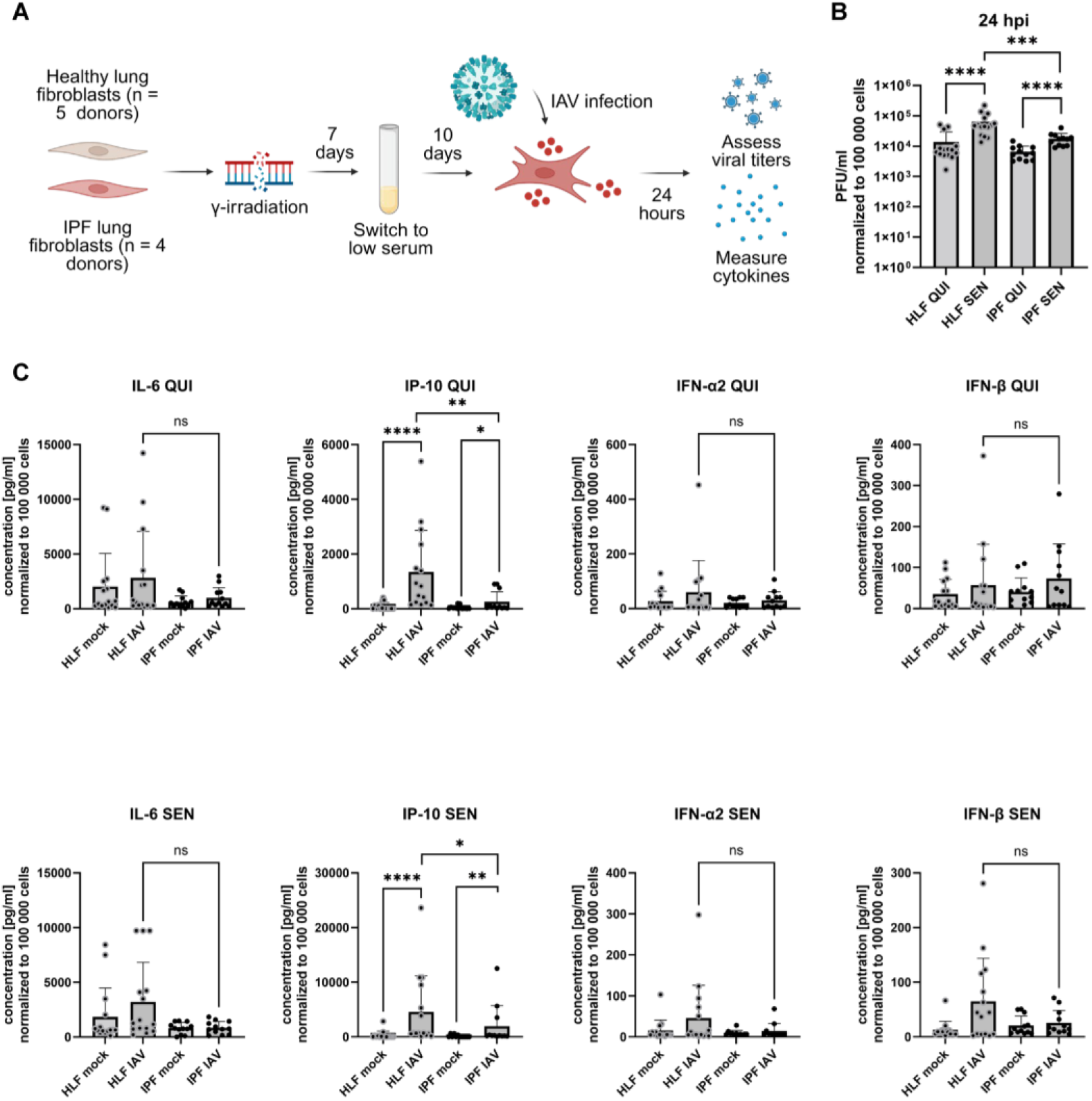
Senescence in IPF fibroblasts results in decreased viral titers and altered cytokine production. (A) Schematic overview of quiescence and senescence induction and subsequent viral infection. (B) Viral titers determined by plaque assay 24 hours post infection (hpi) (healthy lung fibroblasts (HLF): 5 donors with each 3 replicates, idiopathic pulmonary fibrosis (IPF) fibroblasts: 4 donors with each 3 replicates). Plaque-forming units (PFU) have been normalized to cell count and are displayed as PFU/ml adjusted to 100,000 cells. (C) Quantification of cytokines (IL-6, IP-10, IFN-ɑ2, and IFN-β) 24 hpi in the supernatant of mock and infected quiescent (QUI) and senescent (SEN) HLF as well as IPF cells (HLF: 5 donors with each 3 replicates, IPF: 4 donors with each 3 replicates). Cytokine concentration has been normalized to the cell count and is shown as concentration in pg/ml adjusted to 100,000 cells. Statistical significance was calculated using Mann-Whitney U test (B, C). * p ≤ 0.05; ** p ≤ 0.01; **** p ≤ 0.0001.

Overall, our findings suggest that senescence induction in IPF lung fibroblasts results in a transformed reaction to viral infections, characterized by both diminished viral replication and reduced cytokine secretion.

### 2.3 Senescent healthy lung fibroblast gene expression signature is connected to viral responses and the cell cycle

We next wanted to uncover targets driving the lack of functional response to infection in senescent IPF lung fibroblasts. To do this, we used weighted gene correlation network analysis (WGCNA), which generates correlation networks separate from differential expression analysis, to uncover gene and gene networks driving the senescent IPF infected cell state (Figure 3A) (Langfelder & Horvath 2008). The WGCNA revealed two significantly correlated modules, Black and Dimgrey, to the senescent healthy infected condition that were not significantly correlated to the senescent IPF infected condition (Figure 3B). The Black module genes were enriched for genes associated with viral response while the Dimgrey module enriched for genes associated with the cell cycle (Figure 3C). This seems consistent with our differential expression and functional analyses as the senescent IPF lung fibroblasts upregulate more genes related to viral response but fail to functionally respond to a viral infection while senescent healthy lung fibroblasts have a stronger, connected response to viral infection, as shown functionally and by the WGCNA. Individual genes in each module, such as *RIGI* in the Black module, show potential targets driving the senescent normal infected cell state and reinforce our differential expression and functional analyses (Figure 3D).

**Figure 3.**
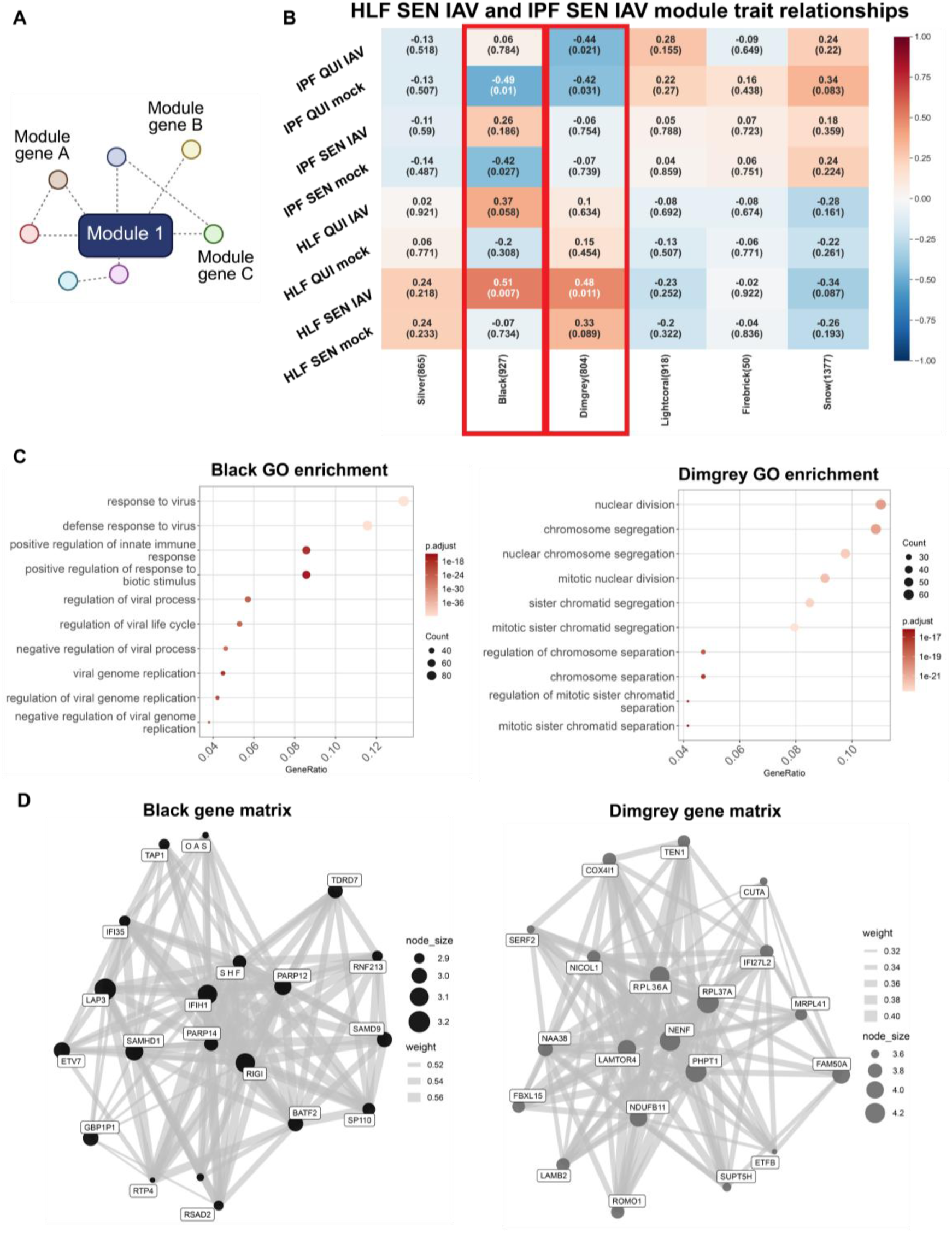
Senescent healthy lung fibroblast gene expression signature is strongly connected to viral responses and the cell cycle. (A) Schematic of WGCNA module connections between genes within a module. (B) WGCNA module-trait heatmap relationships. The rows are the conditions (disease, condition, infection) and each column is a module eigengene. Each cell includes corresponding correlation to the specific module and correlation is indicated by the value in the box. Correlation ranges from -1 to 1 and blue indicates a negative correlation while red indicates a positive correlation. Correlation values are above the number in parenthesis which indicates the p value of each module’s connectivity. Rectangular red boxes are drawn around intersecting rows and columns of interest. HLF = healthy lung fibroblasts. (C) Top 10 GO enrichment terms of genes from the Black (left) and Dimgrey (right) modules. (D) Module gene network visualization for the Black (left) and Dimgrey (right) modules. Nodes represent the top module genes ranked by connectivity z-score, node size scales with intramodular connectivity, and edges represent weighted topological overlap matrix (TOM) connections (TOM ≥ 0.001; top 8 connections retained per gene).

### 2.4 Transcription factor inference identifies differential IRF3/STAT1-associated antiviral programs in senescent healthy and IPF lung fibroblasts

To identify upstream regulators of the distinct antiviral states, we performed transcription factor (TF) enrichment analysis on the RNA-seq data (Müller-Dott et al. 2023). This approach uses perturbation-derived datasets and has been validated in human systems (Müller-Dott et al. 2023). The TF prediction analysis predicted a broadly similar set of virus-associated TFs in senescent healthy and IPF fibroblasts after infection. However, the inferred level of activation differed between the two conditions relative to senescent, mock-infected healthy and IPF lung fibroblasts (Figure 4A). *IRF3* emerged as a prominent candidate and was predicted to be strongly activated in both senescent healthy and IPF fibroblasts after infection (Figure 4A). This was of particular interest because *IRF3* has been less well studied in the context of senescence, fibrosis, and viral infection. *STAT1* was also highlighted because it is a well-established regulator of antiviral interferon responses (Figure 4A). In our analysis, *STAT1* showed differential predicted activation levels between conditions, as *STAT1* in senescent healthy-infected fibroblasts had a TF activity score of between 5-10 while senescent IPF infected fibroblasts had a TF activity score of over 10 (Figure 4A). To support these predictions, we mapped the individual genes contributing to each TF-associated signature. This revealed strong concordance between inferred TF activity and the underlying gene expression patterns in both conditions (Figure 4B).

**Figure 4.**
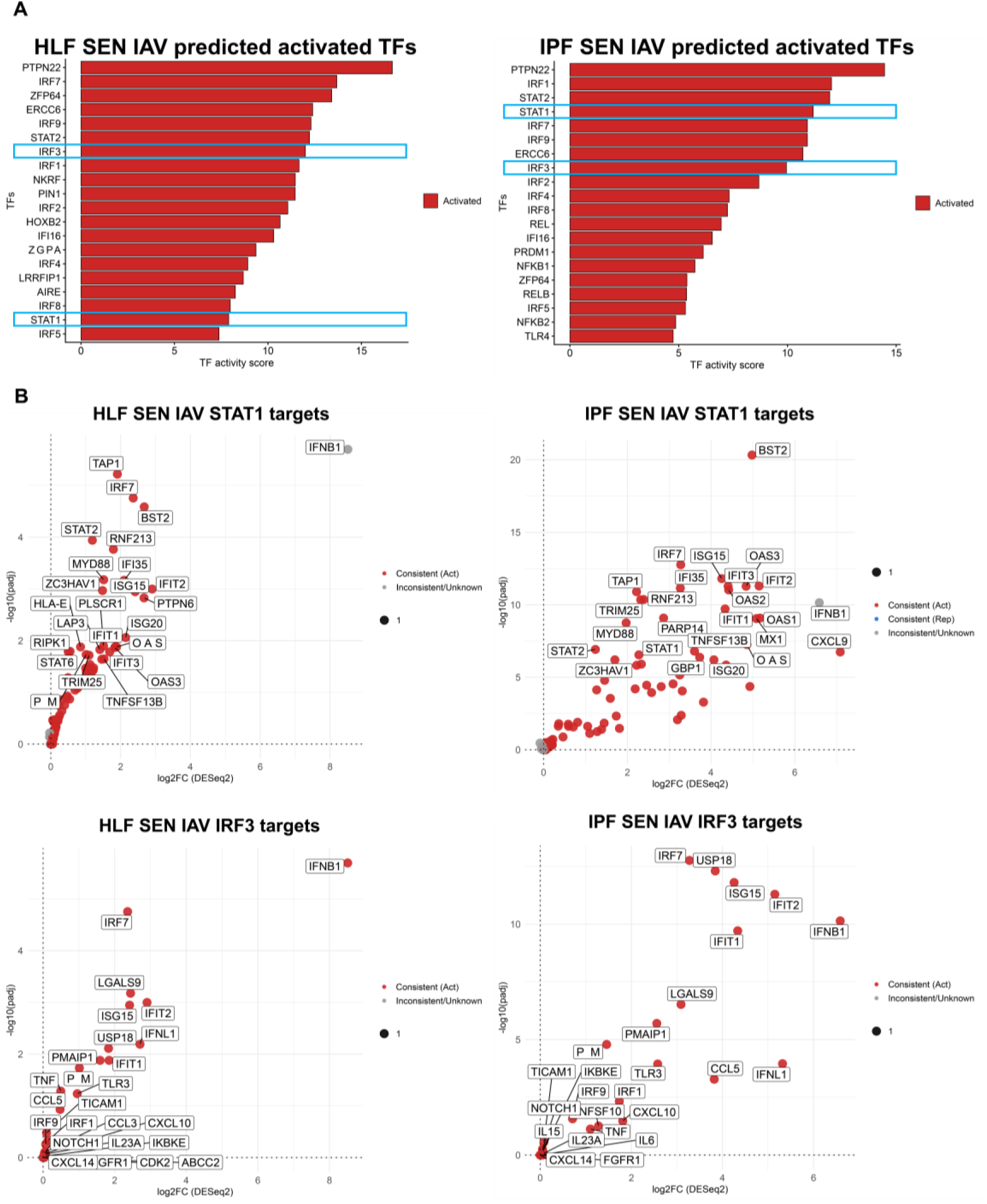
Transcription factor inference identifies differential IRF3/STAT1-associated antiviral programs in senescent healthy and IPF lung fibroblasts. (A) Top activated transcription factors (TF) inferred by decoupleR (ULM) using CollecTRI and DESeq2 shrunken log2FC signatures for healthy (HLF) SEN IAV (left) or IPF SEN IAV lung fibroblasts (right). Bars indicate TF activity scores (red, activated; blue, repressed). No repressed TFs were predicted for either comparison. TFs of interest highlighted in rectangular blue boxes. (B). Healthy SEN IAV (left panels) and IPF SEN IAV (right panels) CollecTRI target genes for *STAT1* and *IRF3*, with points colored by concordance between predicted TF regulation and observed RNA-seq differential expression.

Together, these data suggest that senescent healthy and IPF fibroblasts engage a similar antiviral TF repertoire after infection but differ in the relative activation of key interferon-associated regulators, particularly *IRF3* and *STAT1*.

### 2.5 IRF3 and STAT1 are required for antiviral control in senescent healthy fibroblasts

TF analysis identified *IRF3* and *STAT1* as candidate regulators of the altered antiviral state in infected senescent lung fibroblasts. To test their functional contribution, quiescent and senescent healthy and IPF fibroblasts were transfected with non-targeting (NT), *IRF3*, or *STAT1* siRNA 24 h prior to IAV infection (Figure 5A). Downregulation of mRNA expression level confirmed efficient knockdown of both *IRF3* and *STAT1* across all conditions (Figure 5B), and a modest, non-significant reduction at the protein level was additionally verified in senescent healthy fibroblasts (Supplementary Figure 1A).

**Figure 5:**
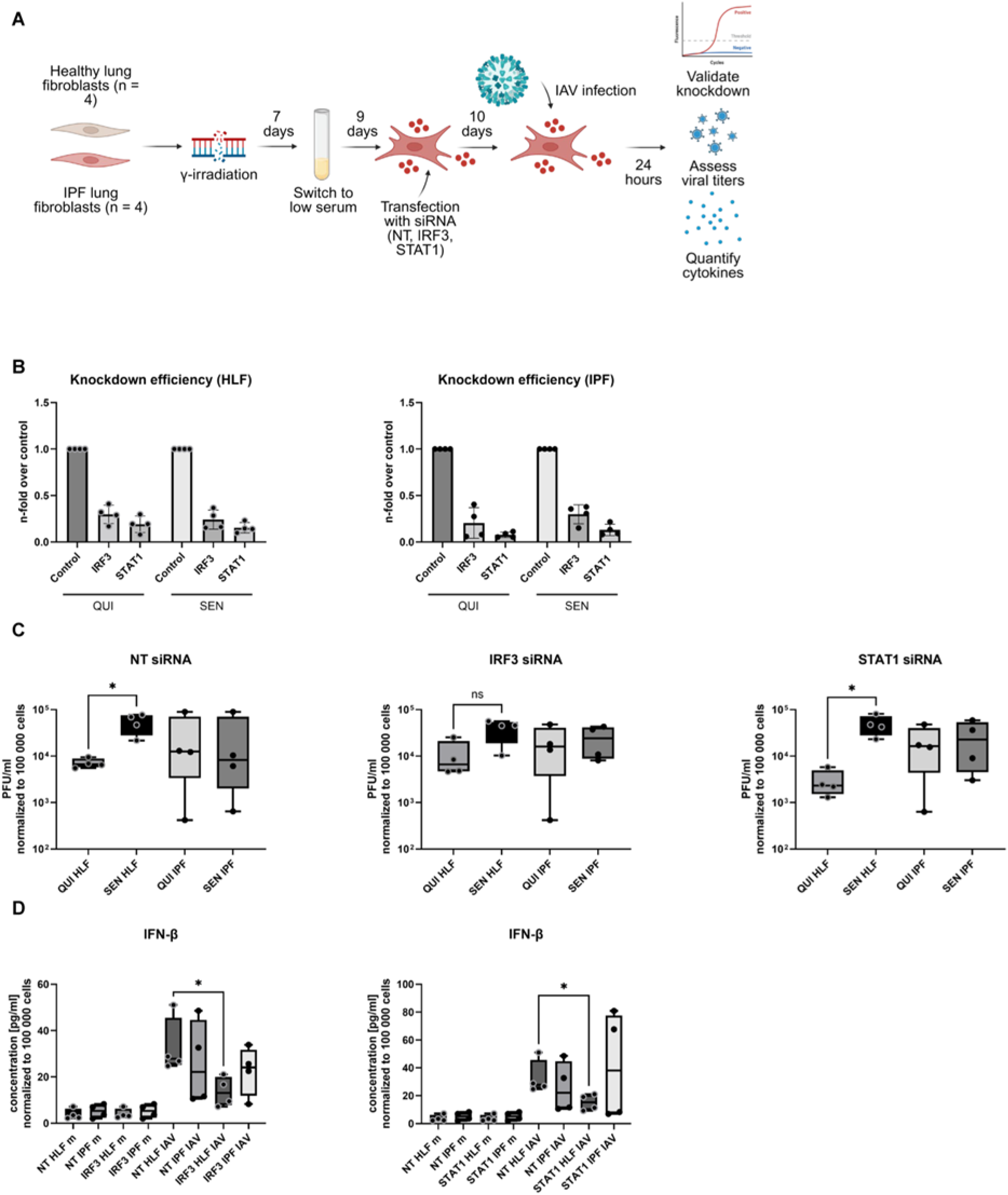
IRF3 and STAT1 are required for antiviral control in senescent healthy fibroblasts. (A) Schematic overview of transfection workflow of quiescent (QUI) and senescent (SEN) healthy lung fibroblasts (HLF) and idiopathic pulmonary fibrosis (IPF) fibroblasts with Accell siRNA (NT, *IRF3* and *STAT1*) 24 hours prior to infection. (B) Assessment of knockdown efficiency for *IRF3* and *STAT1* knockdown on expression level via qRT-PCR (n = 4). Expression is shown as fold change relative to the control NT siRNA condition. (C) Viral titers 24 hours post infection (hpi) in all transfected conditions determined via plaque assay (n = 4). PFU has been normalized to cell count and is displayed as PFU/ml adjusted to 100,000 cells. (D) Cytokine quantification of IFN-β in supernatants from SEN HLF and IPF fibroblasts transfected with NT, *IRF3* (left), or *STAT1* (right) siRNA under mock or infection conditions at 24 hpi. Cytokine concentration has been normalized to cell count and is displayed as concentration in pg/ml adjusted to 100,000 cells. Statistical significance was calculated using Mann-Whitney U test (C, D). * p ≤ 0.05.

Functional perturbation revealed that senescent healthy fibroblasts were partially dependent on *IRF3*/*STAT1* signaling for antiviral control. In these fibroblasts, *IRF3* knockdown was accompanied by a lack of increase in viral plaque formation at 24 hpi (Figure 5C). Additionally, depletion of either *IRF3* or *STAT1* significantly reduced IFN-β secretion after infection in senescent healthy fibroblasts (Figure 5D). In contrast, senescent IPF fibroblasts showed only milder changes following knockdown of *IRF3* or *STAT1*, indicating that the antiviral response of senescent healthy fibroblasts relies more strongly on these signaling nodes than the corresponding response in senescent IPF fibroblasts. Thus, despite retaining transcriptional evidence of antiviral pathway activation, senescent IPF fibroblasts appear more independent of *IRF3*/*STAT1* interferon signaling after infection relative to senescent healthy fibroblasts.

To determine whether siRNA transfection influenced senescence-associated cell state stability, expression of *CDKN1A*, *CDKN2A*, and *LMNB1* was assessed in transfected fibroblasts (Supplementary Figure 1B). Although variability in senescence marker expression was observed after transfection, *CDKN1A* increased and *LMNB1* decreased in senescent healthy fibroblasts with *IRF3* knockdown. In contrast, in senescent IPF fibroblasts, only a significant downregulation of *LMNB1* was identified, which was exclusively observed under the NT condition, indicating alterations of the senescence phenotype in *IRF3*-transfected IPF cells. This data supports the overall functional model in which residual antiviral control in senescent IPF fibroblasts depends less on *IRF3*/*STAT1* signaling after infection relative to senescent healthy fibroblasts.

Together, these findings provide functional validation of the transcription factor inference analysis and identify *IRF3*/*STAT1* as key effectors of antiviral defense in senescent healthy lung fibroblasts that are not functionally activated in IPF lung fibroblasts.

## 3 DISCUSSION

Cellular senescence is increasingly recognized as a central feature of the aging and fibrotic lung, yet its consequences for intrinsic antiviral defense in stromal cells have remained poorly defined. In our study on senescent IPF lung fibroblasts, rather than exhibiting heightened inflammatory activation or generalized dysfunction, senescent IPF fibroblasts adopt an antiviral response state that is transcriptionally but not functionally engaged. Canonical virus-response pathways can still be induced after IAV infection, but their downstream translation into coordinated antiviral control and inflammatory output is altered by the combination of senescence and fibrotic cell identity. This distinction is important because it suggests that fibrotic fibroblasts are not immunologically silent, but instead trapped in a maladaptive response state that is transcriptionally but not functionally active..

One of the central observations is that activation of antiviral transcriptional programs does not necessarily equate to effective antiviral defense. Both senescent healthy and senescent IPF fibroblasts upregulated virus-associated pathways after infection, indicating that elements of pathogen sensing remain intact in both contexts. At the same time, senescent IPF fibroblasts showed a broader set of uniquely induced genes, including *OAS* family members, together with a predicted, stronger coordination of antiviral transcriptional networks. These data support a scenario in which viral sensing is initiated, but the response is not integrated into a coherent protective program. This could also coincide with acute exacerbations driving an already heightened response to virus leading to a detrimental, positive feedback loop and ultimately, functional lung decline. Such a state is consistent with broader aging biology, in which cellular stress responses are present but become uncoupled from initiating the appropriate effector function (Haigis & Yankner 2010; Lavretsky & Newhouse 2012; Vitlic et al. 2014).

Our study shows that senescence increases the viral titers relative to quiescent controls in both healthy and IPF conditions, supporting the idea that senescent cell states can promote viral permissiveness (Schulz et al. 2023). However, senescent IPF fibroblasts displayed an altered inflammatory profile, including reduced *CXCL10*/IP-10 secretion, despite retaining transcriptional evidence of antiviral pathway activation. This dissociation between transcriptional induction and inflammatory output suggests that fibrotic fibroblasts fail to properly convert viral recognition into an appropriately coordinated response. In the context of the lung, this has potentially broad implications. Fibroblasts are now more commonly known to be immunologically competent stromal cells capable of sensing danger and influencing epithelial and immune-cell behavior (Cavagnero & Gallo 2022; Ghonim et al. 2023). A dysfunctional antiviral fibroblast state could then contribute to altered host defense and heightened susceptibility to infection-associated deterioration.

The network-level analyses build on this model further. Viral-response and cell-cycle modules were specifically linked to the senescent healthy infected state, whereas comparable coordinated programs were weak or absent in senescent IPF fibroblasts. This finding suggests that fibrotic identity destabilizes the response to viral challenge under senescent conditions. Efficient antiviral defense requires synchronized coupling of sensing, interferon induction, and downstream effector programs. Once compromised, the resulting state may be observed as “antiviral” at the transcript level but fail at the functional level, which requires systems level coordination to enact a functional response from transcriptional signals. The WGCNA results provide an important systems level view between the transcriptomic data and functional phenotype. Although virus-associated genes are activated in senescent IPF lung fibroblasts after infection, the response is uncoordinated within gene networks based on the WGCNA results. This is then associated with a lack of a functional response to viral infection in senescent IPF lung fibroblasts and ultimately supports the view that senescent IPF fibroblasts occupy a partially activated yet biologically incompetent antiviral state.

*IRF3* and *STAT1* emerged as key regulatory nodes within this altered antiviral landscape. Although both transcription factors are central mediators of interferon signaling and were predicted to be activated in infected fibroblasts, their functional relevance differed by cellular context. In senescent IPF fibroblasts, poor coordination of the broader antiviral network may increase reliance on a limited number of regulatory nodes, rendering antiviral control particularly vulnerable to disruption of specific signaling axes such as *IRF3*/*STAT1*-associated signaling. This continues to support the idea that although virus-associated TFs are predicted to be activated in senescent IPF fibroblasts, the appropriate effector function is not initiated. Validating which TFs are functionally active or repressed in senescent IPF fibroblasts would help to understand at which point senescent IPF viral response capabilities could be restored.

This model offers a plausible explanation for the clinical vulnerability of patients with IPF to respiratory viral infections. Acute exacerbations are among the most devastating complications of IPF, and infections are widely considered important triggers or amplifiers of these episodes (Ryerson et al. 2015). A fibroblast compartment that retains partial viral sensing yet fails to mount a coordinated interferon-driven response may contribute to disease worsening by blunting paracrine immune signaling and disturbing the balance between injury and repair in an already damaged lung. The use of primary human healthy donor and patient-derived IPF fibroblasts further strengthens the clinical relevance of this concept.

Overall, cellular senescence does not simply intensify pre-existing fibrotic programs. Rather, the two states show a distinct form of stromal antiviral dysfunction in which canonical sensing pathways remain partly inducible, but their downstream execution is poorly coordinated and functionally fragile. This view places fibroblasts more centrally into the biology of infection susceptibility in the aging fibrotic lung and suggests that therapeutic strategies aimed at restoring coordinated interferon responsiveness may be more relevant than indiscriminately amplifying inflammation.

Taken together, these findings define senescent IPF fibroblasts as a dysfunctional antiviral stromal state shaped by the convergence of cellular senescence and fibrotic cell identity. Preserved elements of viral sensing coexist with impaired systems-level coordination and altered dependency on *IRF3*/*STAT1*-associated interferon signaling. This provides a mechanistic framework linking fibrotic remodeling to impaired local antiviral defense and may help explain why the aging fibrotic lung is particularly vulnerable to virus-associated acute exacerbations and disease worsening.

In conclusion, our findings define senescent IPF fibroblasts as a dysfunctional antiviral fibroblast state characterized by preserved viral sensing but impaired interferon response. By identifying *IRF3*/*STAT1*-associated signaling as a critical, missing control axis in the context of IPF lung fibroblasts, this work also highlights a potential point of therapeutic approaches in the aging fibrotic lung.

## 4 EXPERIMENTAL PROCEDURES

### 4.1 Virus strain and propagation

Viral infections were carried out with Influenza A virus/H1N1/Puerto Rico/8/1934 (IAV), which has been propagated in Madin-Darby canine kidney (MDCK) cells cultivated in minimum essential medium (MEM) containing 10% fetal bovine serum (FBS) and 1% Penicillin/Streptomycin (P/S) (Hornung et al. 2025). Viral titers from both propagated virus and *in vitro* infections were quantified as plaque-forming units (PFU) per milliliter using a plaque assay, as previously detailed (Deinhardt-Emmer et al. 2021). For viral titers obtained from *in vitro* experiments, PFU counts were normalized to cell counts and expressed as PFU per 100,000 cells.

### 4.2 Cultivation of primary human lung fibroblasts from healthy and IPF donors

Healthy lung fibroblasts (HLF) and lung fibroblasts from donors diagnosed with idiopathic pulmonary fibrosis (IPF) were obtained from adult donors from collaborators at the University of California, San Francisco, or Cedars-Sinai and isolated at their originating institutions through methods previously described (Hogaboam et al. 2005). A complete list of details for each donor lung fibroblast can be found in “Supplementary Table 1 - Donor Information.” Cells were cultured in Dulbecco’s Modified Eagle Medium (DMEM) supplemented with 10% FBS and 1% P/S at 37°C with 5% CO_2_.

### 4.3 Induction of cellular senescence

Senescence in lung fibroblasts was induced by irradiation, in which the cells were exposed to a dose of 15 Gray while maintained in cell culture plates. Immediately after irradiation, the medium was replaced with fresh DMEM containing 10% FCS and 1% P/S. Subsequently, the medium was refreshed every 2-3 days. The induction of senescence was completed after 10 days of culture, and the senescent cells were then utilized for further experiments.

As a control, quiescent cells were generated by serum starvation as previously described (Schulz et al. 2023; Neri et al. 2021).

### 4.4 *In vitro* infections

HLF and IPF cells were infected with IAV at a multiplicity of infection (MOI) of 1 in Dulbecco’s Phosphate Buffered Saline (DPBS) containing 0.2% bovine serum albumin (BSA; Roth, Karlsruhe, Germany), 1 mM MgCl_2_ (Sigma Aldrich), and 0.9 mM CaCl_2_ (Sigma Aldrich) for a duration of 30 minutes (min). Following this incubation, the cells were washed and subsequently cultured in DMEM supplemented with 0.2% BSA, 1 mM MgCl_2_, 0.9 mM CaCl_2_, and 30 ng/ml of TPCK-treated trypsin (Thermo Fisher Scientific, Waltham, MA, USA) for 24 hours (h) (Schulz et al. 2023).

### 4.5 RNA Extraction

Cells were first lysed with RLT buffer before RNA extraction was conducted using the RNeasy Mini Kit (Qiagen, Hilden, Germany) according to the manufacturer’s instructions. Concentration and purity of extracted RNA was assessed with the NanoDrop spectrophotometer ND-1000 (Thermo Fisher Scientific/ PeqLab).

### 4.6 mRNA sequencing

Library construction and total mRNA sequencing of quiescent as well as senescent mock and infected HLF (each n=3) and IPF cells (each n=4) was carried out by Novogene Co., LTD (Munich, Germany).

### 4.7 cDNA Synthesis and qRT-PCR

RNA was reverse transcribed to cDNA using the High-Capacity cDNA Kit (Thermo Fisher Scientific). qRT-PCR was subsequently performed with the Maxima SYBR Green qPCR Master Mix (Thermo Fisher Scientific) on a Rotor Gene Q (Qiagen) as previously described(Schulz et al. 2023). Primers were purchased from metabion (Planegg, Germany) and sequences are detailed in “Supplementary Table 2.” Genes of interest were normalized to β-Actin and the -2ΔΔCT method was employed to quantify relative gene expression.

### 4.8 Cytokine quantification

Multiplex cytokine quantification in the supernatant of HLF and IPF fibroblasts was performed using the LEGENDplex™ Hu Anti-Virus Response Panel 1 V02 (Biolegend, San Diego, CA, USA) according to the manufacturers protocol. Measurements were conducted with the FACS Symphony A1 (BD, Franklin Lakes, NJ, USA) and data was analyzed via the Legendplex analysis software Qognit. The concentration of cytokines was normalized to the cell count and adjusted to 100 000 cells.

### 4.9 Transfection with Accell siRNA

Quiescent and senescent HLF and IPF cells were transfected 24 h prior to infection. To successfully knock down the target genes *IRF3* and *STAT1*, a method combining the transfection reagent DharmaFECT3 (Dharmacon, Lafayette, CO, USA) and Accell siRNA (Dharmacon) was adapted(Dong et al. 2020). For siRNA transfection, quiescent and senescent HLF and IPF cells were cultured in DMEM without supplements for 2 h before transfection. A siRNA pool consisting of four individual siRNAs targeting either *IRF3* or *STAT1*, along with a non-targeting control (NT) siRNA, was utilized (*IRF3*: E-006875-00-0020; *STAT1*: E-003543-00-0020; NT: D-001910-10-50; all Dharmacon). siRNAs were diluted in Accell delivery medium to a final concentration of 40 nM per well and were incubated at room temperature (RT) for 5 min. Simultaneously, DharmaFECT 3 transfection reagent was diluted in Accell delivery medium (0.5 µl for 24-well plates or 1 µl for 12-well plates) and also incubated for 5 min at RT. The diluted siRNA and DharmaFECT 3 solutions were then combined at a 1:1 ratio and further incubated at RT for an additional 20 min.

After adding 100 µl to 24-well plates or 200 µl to 12-well plates, the transfected cells were cultivated for 24 h at 37°C in 5% CO_2_. Transfection efficiency was assessed by qRT-PCR to measure the reduction in target gene expression.

### 4.10 Bulk RNA-seq processing

Raw reads were aligned locally using the nf-core/rnaseq salmon pipeline in windows system linux, performing pseudo-alignment against a standard reference transcriptome for Homo sapiens (GRCh38 genome) (Ewels et al. 2020; Patro et al. 2017). Transcript-level count estimates (estimated counts and TPMs) per sample were then summarized to gene-level count matrices using the corresponding transcript-to-gene annotation in R. The gene-level estimated count matrices along with sample metadata were used for bulk RNA-seq dataset generation and analyses. Sample metadata can be found in “Supplementary Table 3 - Jena_Buck_IAV_metadata.”

### 4.11 Differential expression analysis

DESeq2 formula used is provided here for the lung fibroblasts: ∼ disease * infection * condition (Love et al. 2014). Principal component analysis (PCA) provided in Figure 1B showing rationale for DESeq2 formula. GO and KEGG enrichment performed as previously described (Consortium et al. 2025; Ashburner et al. 2000; Kanehisa & Goto 2000). Transcription factor (TF) enrichment was performed using the Collection of Transcription Regulation Interactions meta-resource that was recently sign curated for activation or repression of each TF in the database (Müller-Dott et al. 2023). Differentially expressed genes (DEG) lists can be found in “DEG_normal_IR_IAV_vs_normal_IR_mock.csv” and “DEG_IPF_IR_IAV_vs_IPF_IR_mock.csv.”

### 4.12 Weighted gene correlation network analysis (WGCNA)

WGCNA was run using pyWGCNA (Rezaie et al. 2023). The topological overlap matrix similarity dendrogram was made using the R WGCNA package and the gene matrices were made using the BioMart R package (Langfelder & Horvath 2008). The gene expression matrix used for pyWGCNA was batch-corrected and ran through a variance stabilizing transformation which was then filtered for top 75% most varying genes. Resulting gene modules were tested for gene set enrichment analysis (GSEA) against Gene Ontology (GO) Terms, as previously described, and cross-referenced with DEGs (Ashburner et al. 2000). Our total sample number did not reach the commonly recommended sample number for sufficient power, which is 15 total samples, but meaningful results can still be discovered with less than 15 samples and emphasizes the importance of validating WGCNA module genes regardless of sample size (Langfelder & Horvath 2008; Silva et al. 2022). WGCNA module gene and DEG overlap lists provided in “_overlap.csv” tables.

### 4.13 Western blot

For protein extraction, cells were lysed with RIPA buffer supplemented with Halt Protease and Phosphatase Inhibitor (Thermo Fisher Scientific), followed by protein quantification via Micro BCA Protein Assay Kit (Thermo Fisher Scientific). Protein separation was conducted on a 12% SDS-polyacrylamide gel at 30 mA for 60 min, followed by a transfer to polyvinylidene fluoride membranes at 150 mA for 2 h. Membranes were blocked for 1 h at RT with 5% milk-Tris-buffered saline with Tween-20 (TBST) and then incubated overnight at 4°C with primary antibodies (IRF3 (D83B9) Rabbit Monoclonal Antibody and STAT1 Antibody, both 1:1000 dilution, Cell Signaling Technology, Danvers, MA, USA).

Secondary HRP-conjugated antibodies were diluted 1:1000 in 5% milk-TBST and incubated for 1 h at room temperature. β-Actin and HRP-Vinculin were utilized as housekeeping proteins. Membrane images were captured using the iBright™ CL750 Imaging System (Thermo Fisher Scientific) and quantified with ImageJ. Protein concentrations were expressed relative to the respective NT control.

### 4.14 Statistical analysis and Illustrations

Statistical analyses were conducted using GraphPad Prism 10. The specific statistical tests employed are detailed within the corresponding figure legends. Illustrations were created with the aid of <u>Biorender.com</u>.

### 4.15 Code availability

Code will be made available on github upon acceptance of manuscript.

## Supporting information

20260415_Buck Jena paper_supplemental

DEG_IPF_IR_IAV_vs_IPF_IR_mock

DEG_IPF_IR_IAV_vs_IPF_IR_mock__black__overlap

DEG_IPF_IR_IAV_vs_IPF_IR_mock__dimgrey__overlap

DEG_normal_IR_IAV_vs_normal_IR_mock

DEG_normal_IR_IAV_vs_normal_IR_mock__black__overlap

DEG_normal_IR_IAV_vs_normal_IR_mock__dimgrey__overlap

Supplementary Table 1 - Donor Information

Supplementary Table 3 - Jena_Buck_IAV_metadata

## DECLARATIONS

### Author Contributions

Conceptualization: J.W.B., P.Y.D., S.D.-E.; methodology: F.H., Y.R., J.W.B.; Cell provision: J.W.B., A.L.C., T.H., C.J.L.S.; bioinformatics: J.W.B., T.A.U.H., F.S. K.S.; writing—original draft preparation: J.W.B., P.Y.D., S.D.-E., Y.R.; writing—review and editing, all authors; visualization: J.W.B., P.Y.D., S.D.-E., Y.R.; resources and supervision: S.D.-E., P.Y.D., D.F., C.M.H., C.J.L.S.; funding acquisition, S.D.-E., P.Y.D. All authors have read and agreed to the published version of the manuscript.

## Acknowledgements

We would like to acknowledge Dr. Judith Campisi and the legacy she built in the field of senescence that we hope to continue through this paper. We would like to thank the Buck Institute for letting the lab continue until we finish publishing papers started by Judy. For their exceptional technical contributions, we would also like to thank Lea Herrmann, Yvonne Ozegowski and Monique Keller. ChatGPT (https://chatgpt.com/) was used for the preparation of the manuscript to improve writing in terms of phrasing, grammar and spelling. The content was then evaluated by the authors and adjusted as necessary.

## Data availability statement

Raw sequencing data (FASTQ files) and processed bulk RNA-seq data generated in this study will be deposited in the Gene Expression Omnibus/ArrayExpress (GEO); accession numbers will be provided upon acceptance of the manuscript. Combined metadata can be found in “Table - Jena_Buck_IAV_metadata.” Additional data that support the findings of this study are available from the corresponding author upon reasonable request.

