## Supplementary material for "Cellular senescence dysregulates antiviral interferon responses in idiopathic pulmonary fibrosis": 20260415_Buck Jena paper_supplemental

### Supplementary Figure 1

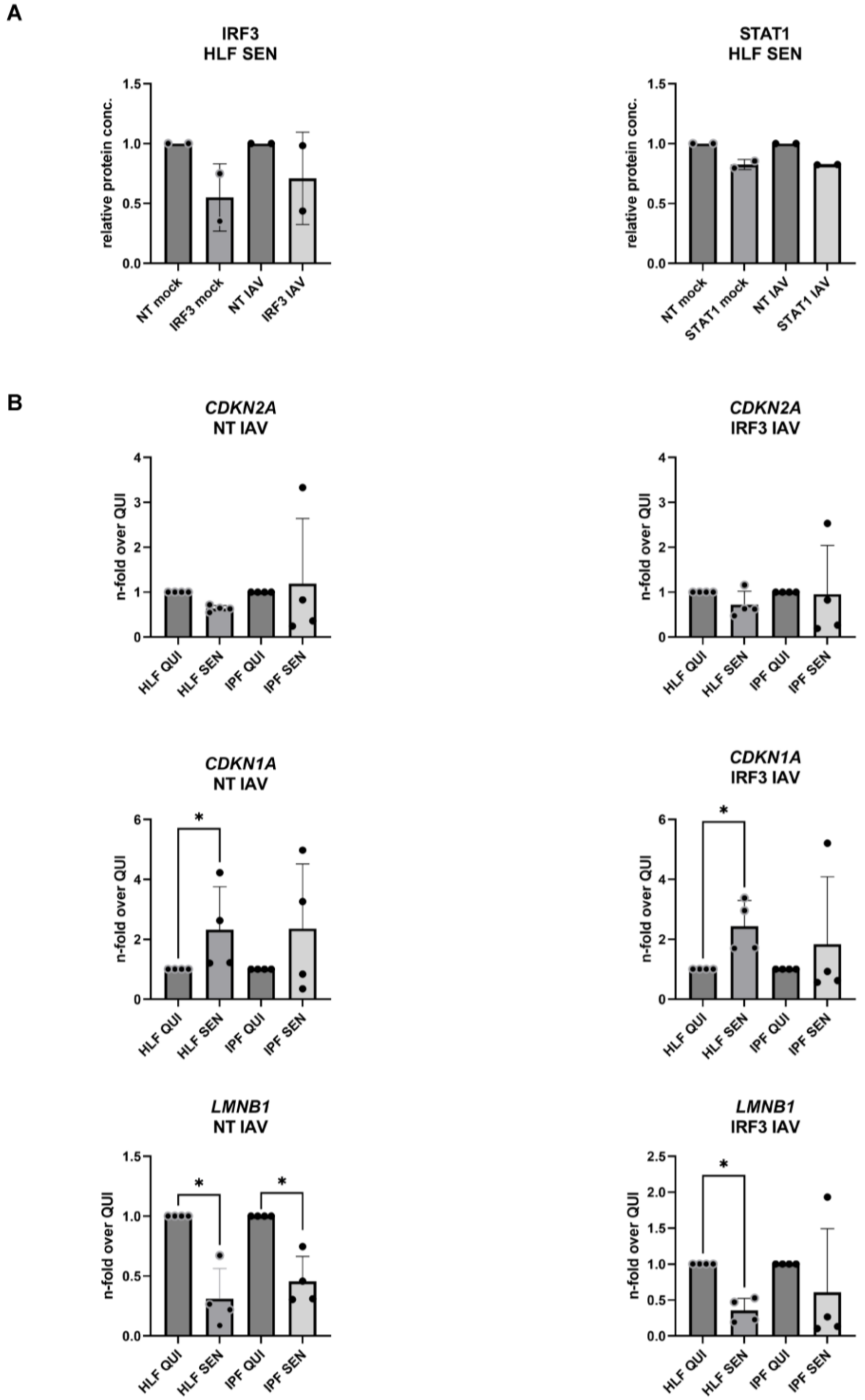

(A) Quantification of *IRF3* and *STAT1* proteins in senescent (SEN) HLF healthy lung fibroblasts via Western blot (n = 2). Protein levels were quantified using Image J and are normalized to the respective housekeeping protein (Vinculin for *IRF3* and β-Actin for *STAT1*) and are expressed relative to the respective NT control siRNA condition. (B) The relative expression of senescence markers *CDKN1A* (p21), *CDKN2A* (p16), and *LMNB1* (Lamin B1) was quantified by qRT-PCR in both infected quiescent (QUI) and SEN HLF and idiopathic pulmonary fibrosis (IPF) fibroblasts transfected with NT or *IRF3* siRNA (n = 4). Gene expression levels were normalized to β-Actin and reported relative to the respective quiescent control. Statistical significance was determined using the Mann-Whitney U test (B). * $p \leq 0.05$.

**Supplementary Table 2**

| **Gene name** | **Forward sequence** | **Reverse sequence** |
|---|---|---|
| *IRF3* | TCTGCCCTCAACCGCAAAGAAG | TACTGCCTCCACCATTGGTGTC |
| *STAT1* | ATGGCAGTCTGGCGGCTGAATT | CCAAACCAGGCTGGCACAATTG |
| *ACTB* | CATGTACGTTGCTATCCAGGC | CTCCTTAATGTCACGCACGAT |
| *LMNB1* | TTGGATGCTCTTGGGGTTC | AAGCAGCTGGAGTGGTTGTT |
| *CDKN1A* (p21) | TCACTGTCTTGTACCCTTGTGC | GGCGTTTGGAGTGGTAGAAA |
| *CDKN2A* (p16) | CTCGTGCTGATGCTACTGAGGA | GGTCGGCGCAGTTGGGCTCC |

Forward and reverse primer sequences used in qRT-PCR. All primers are human.
